# METTL3 promotes homologous recombination repair and modulates chemotherapeutic response by regulating the EGF/Rad51 axis

**DOI:** 10.1101/2021.11.11.468249

**Authors:** Enjie Li, Mingyue Xia, Yu Du, Kaili Long, Feng Ji, Feiyan Pan, Lingfeng He, Zhigang Hu, Zhigang Guo

## Abstract

METTL3 and N6-methyladenosine (m6A) are involved in many types of biological and pathological processes, including DNA repair. However, the function and mechanism of METTL3 in DNA repair and chemotherapeutic response remain largely unknown. In present study, we identified that METTL3 participates in the regulation of homologous recombination repair (HR), which further influences chemotherapeutic response in breast cancer (BC) cells. Knockdown of METTL3 sensitized BC cells to Adriamycin (ADR) treatment and increased accumulation of DNA damage. Mechanically, we demonstrated that inhibition of METTL3 impaired HR efficiency and increased ADR-induced DNA damage by regulating m6A modification of EGF/RAD51 axis. METTL3 promoted EGF expression through m6A modification, which further upregulated RAD51 expression, resulting in enhanced HR activity. We further demonstrated that the m6A “reader,” YTHDC1, bound to the m6A modified EGF transcript and promoted EGF synthesis, which enhanced HR and cell survival during ADR treatment. Our findings reveal a pivotal mechanism of METTL3-mediated HR and chemotherapeutic drug response, which may contribute to cancer therapy.

## Introduction

N^6^-methyladenosine (m^6^A) modification of RNA has been reported to participate in regulating numerous cellular processes in eukaryotes (Wang et al., 2014; Wang et al., 2015; Xiang et al., 2017). Methyltransferase-like 3 (METTL3) is a key member of the m^6^A methyltransferase complex, which also includes co-factors METTL14 and the Wilms tumor 1 associated protein (WTAP) (Liu et al., 2014; Wang *et al*., 2015). This RNA modification can be recognized by a set of m^6^A-binding proteins, including YTH domain containing family protein (YTHDF1/2/3), YTH domain containing 1/2 (YTHDC1/2), and insulin like growth factor 2 mRNA binding protein (IGF2BP1/2/3), which serve as m^6^A “readers” and mediate specific functions of m^6^A-modified RNA (Deng et al., 2018; Wang *et al*., 2015). Furthermore, fat mass and obesity-associated protein (FTO) and RNA demethylase, ALKBH5, work as m^6^A “erasers” to remove m^6^A modifications from RNA (Deng *et al*., 2018). Recent studies have indicated that m6A modification and METTL3 play important roles in the progression and chemotherapy response of various cancers, including BC (Cai et al., 2018; Deng *et al*., 2018; Pan et al., 2021).

Chemotherapy is used in early-stage BC and locally advanced BC to provide an improved chance for breast-conserving surgery, reduce the risk of recurrence, and increase survival rates (Fisusi and Akala, 2019). The use of adjuvant chemotherapy remains an effective treatment for triple-negative BC and other types of invasive breast cancer (Hennigs et al., 2016). Several chemotherapeutic agents, including ADR, docetaxel, 5-fluorouracil, and cisplatin are used in combination chemotherapy for BC treatment (Fisusi and Akala, 2019). Chemotherapeutic agents, such as ADR, induce apoptosis by causing DNA damage (He et al., 2016; Lu et al., 2020). Targeting the DNA damage response (DDR) may enhance the sensitization of cancer cells to chemotherapeutic drugs (Li et al., 2021; Lu *et al*., 2020). Moreover, elevated DNA repair activity contributes to the drug resistance of cancer cells treated with chemotherapy (Li *et al*., 2021; Lu *et al*., 2020).

Key proteins involved in DNA repair, such as BRCA1 and Rad51 in homologous recombination repair (HR), xeroderma pigmentosum group C (XPC) in nucleotide excision repair (NER), and flap endonuclease 1 (FEN1) in base excision repair (BER), influence the susceptibility of various cancers and may be suitable targets in cancer mono- and combination therapy (Ali et al., 2017; Grundy et al., 2020; Li *et al*., 2021; Lu *et al*., 2020; Malik et al., 2020; Miki et al., 1994). However, targeting key DDR proteins to improve sensitization of cancer cells and circumvent cancer cell resistance remain significant challenges to achieving satisfactory therapeutic effects (Li *et al*., 2021; Pan *et al*., 2021). Therefore, exploring novel treatment strategies and mechanisms that affect DDR in cancer cells may contribute to improved treatment for these diseases.

Currently, METTL3 and m^6^A modification have been implicated in DDR (Xiang *et al*., 2017; Yu et al., 2021b; Zhang et al., 2020). However, the molecular mechanism of METTL3 and m^6^A modification in DDR, especially in the context of chemotherapeutic drug-induced DDR, remains unexplored. Here, we report that METTL3 is involved in the regulation of HR by regulating the EGF/RAD51 axis. Knockdown of METTL3 sensitizes BC cells to ADR, impairs HR, and induces significant DNA damage. Moreover, YTHDC1 was identified as the reader that binds and protects m^6^A-modified EGF mRNA and regulates DNA repair and the response of BC cells to ADR. Overall, our findings provide insight into the function and mechanism of METTL3 in HR and the chemotherapeutic drug response in BC, and demonstrate the potential of targeting METTL3 as an antitumor treatment.

## Results

### METTL3 regulates chemotherapeutic response of BC cells

METTL3 has been reported to be involved in the progression of several types of cancers, including BC (Deng *et al*., 2018; Wang et al., 2020a). We wonder whether METTL3 regulates chemotherapeutic response of BC cells. First, we identified the elevated METTL3 and m^6^A levels in BC cells (Supplementary Fig. S1A, B). Then, we investigated the role of METTL3 in regulating the sensitivity of BC cells to chemotherapeutic drugs with stable METTL3-OV or -KD cell lines (Supplementary Fig. S1C, D). Cell viability assays were done using METTL3-KD and METTL3-OV MCF-7 stable cell lines treated with five first-line chemotherapeutic drugs including 5-FU, cisplatin (DDP), Adriamycin (ADR), paclitaxel, and carboplatin. The results indicated that modification of METTL3 expression markedly attenuated the sensitivity of MCF-7 cells to ADR compared with the other drugs (Fig. 1A and Supplementary Fig. S1E-I) (Andreetta et al., 2010). Cell viability assays using METTL3-OV and -KD MB-231 stable cells verified the effect of METTL3 on ADR sensitivity (Fig. 1B and supplementary Fig. S1E). These results were further verified in a morphological analysis which showed that silencing METTL3 enhanced chemotherapeutic drug sensitivity (Fig. 1C), whereas overexpression of METTL3 decreased sensitivity in MCF-7 cells (Supplementary Fig. S1J). Furthermore, flow cytometry analysis demonstrated that treatment with Adriamycin induced higher apoptosis rates in both METTL3-KD MCF-7 and MB231 cells compared with control cells (Fig. 1D and Supplementary Fig. S1K). Accordingly, elevated pro-apoptotic Bax and caspase 3 were detected in METTL3-KD BC cells treated with Adriamycin (Fig. 1E). As expected, silence of METTL3 reduced global m^6^A levels in BC cells, whereas overexpression of METTL3 increased m^6^A levels in MCF-7 and MB-231 cells compared with control cells (Supplementary Fig. 1L, M).

**Figure 1.**
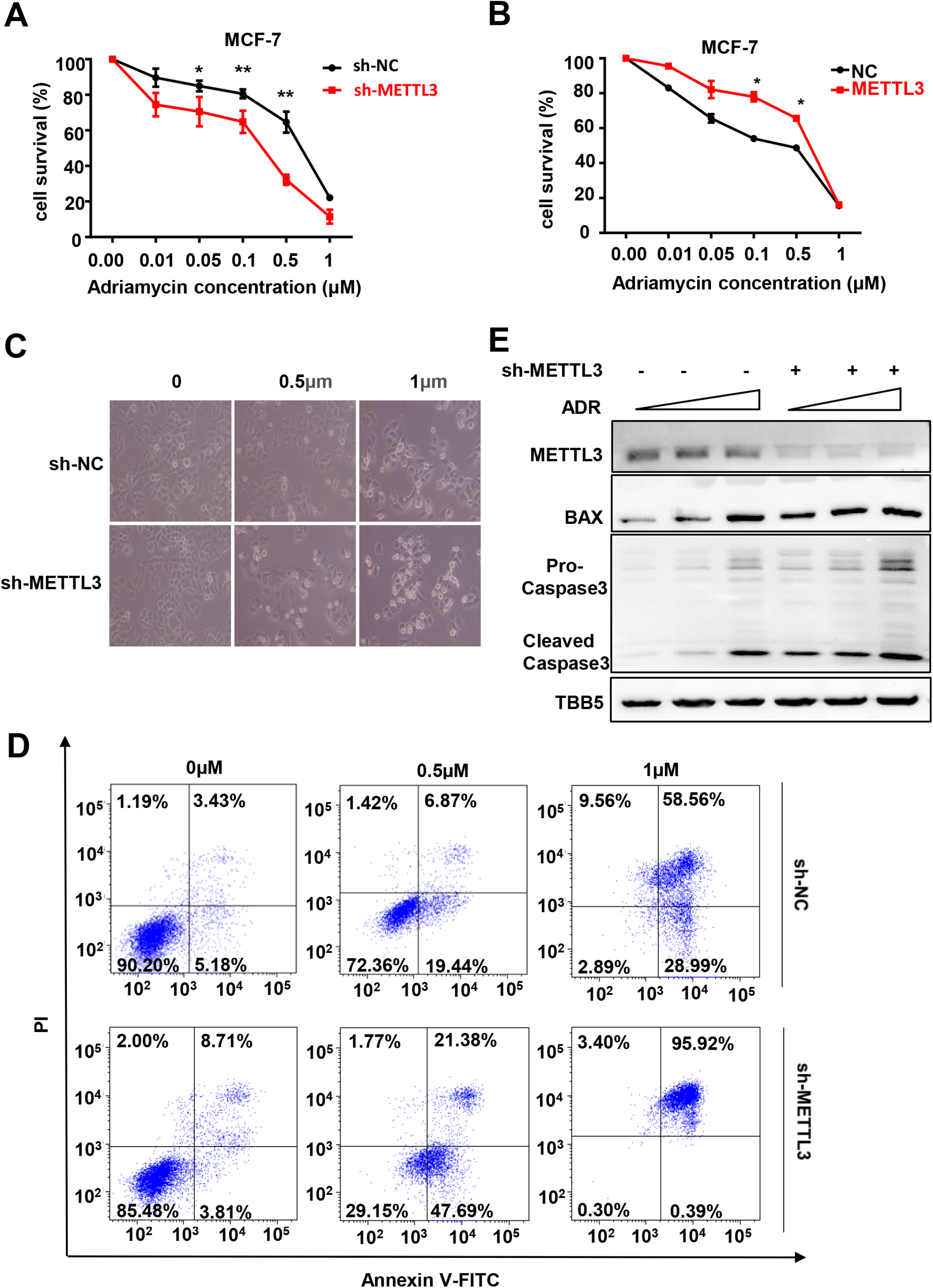
Knockdown of METTL3 sensitizes BC cells to ADR. (**A, B**) MTT assays were performed to determine the effect of METTL3 on ADR cytotoxicity in MCF-7 or MDA-MB231 cells. Data are expressed as the mean ± standard deviation (SD), n=3 per group. (**C**) Morphological analysis of MCF-7 with different drug treatments. (**D**) Annexin V/PI staining and flow cytometry assay of control or METTL3-KD MCF-7 cells with different drug treatments. (**E**) Western blot analysis of METTL3, BAX, and Caspase 3 in control or METTL3-KD MCF-7 cells treated with various concentrations of ADR. ^*^ P< 0.05; ^**^ P< 0.01; ^***^ P< 0.001.

### METTL3 promotes HR

Adriamycin is normally described as a classic topoisomerase II poison that intercalates into DNA and forms DNA adducts, and subsequently induces DNA double strand breaks (DSBs) (Swift et al., 2006; Yang et al., 2014). We further addressed whether METTL3 was involved in DSB repair and affected ADR-induced DNA damage. To explore the role of METTL3 in the regulation of DNA repair, we treated METTL3-KD or –OV BC cells with ADR and subsequently released the cells into fresh medium lacking ADR and monitored the levels of γH2AX (an established marker of DNA damage) over time. Our data showed that knockdown of METTL3 maintained higher γH2AX levels in both MCF-7 and MB-231 cells compared with control cells (Fig. 2A, B), whereas overexpression of METTL3 resulted in an earlier decline of γH2AX compared with control cells (Supplementary Fig. S2A, B). We further detected the effect of METTL3 on the regulation of DSB repair that was induced by etoposide (ETO; another inhibitor of topoisomerase II). Accordingly, our data showed that METTL3 promoted the repair of DSB induced by ETO in both MCF-7 and MB-231 cells (Fig. 2C, D; and Supplementary Fig. S2C, D). Consistently, an increased number of positive nuclei foci of γH2AX was detected in both METTL3-KD MCF-7 and MB-231 cells after ADR treatment and then released after four hours (Fig. 2E, F), whereas experiments using METTL3-OV BC cells showed the opposite effects (Supplementary Fig. S2E). Similar results were obtained for the foci of 53BP1 (tumor-suppressor p53-binding protein 1, a key regulator of DSB repair) in METTL3-KD and METTL3-overexpressing MCF-7 cells (Supplementary Fig. S2F). DSBs are primarily repaired by either HR or non-homologous end joining (NHEJ) (Sonoda et al., 2006). Using two well-characterized green fluorescent protein (GFP)-based HR and NHEJ reporter systems, we determined the effect of METTL3 on HR and NHEJ efficiency (Fig. 2G and Supplementary Fig. S2G). The results showed that overexpression of METTL3 significantly enhanced HR-mediated DSB repair (Fig. 2H), whereas no effect was observed on NHEJ-mediated DSB repair (Supplementary Fig. S2H). This is consistent with previous reports using the HR and NHEJ luciferase reporter system based on crisper-cas9-induced DSBs (Zhang *et al*., 2020).

**Figure 2.**
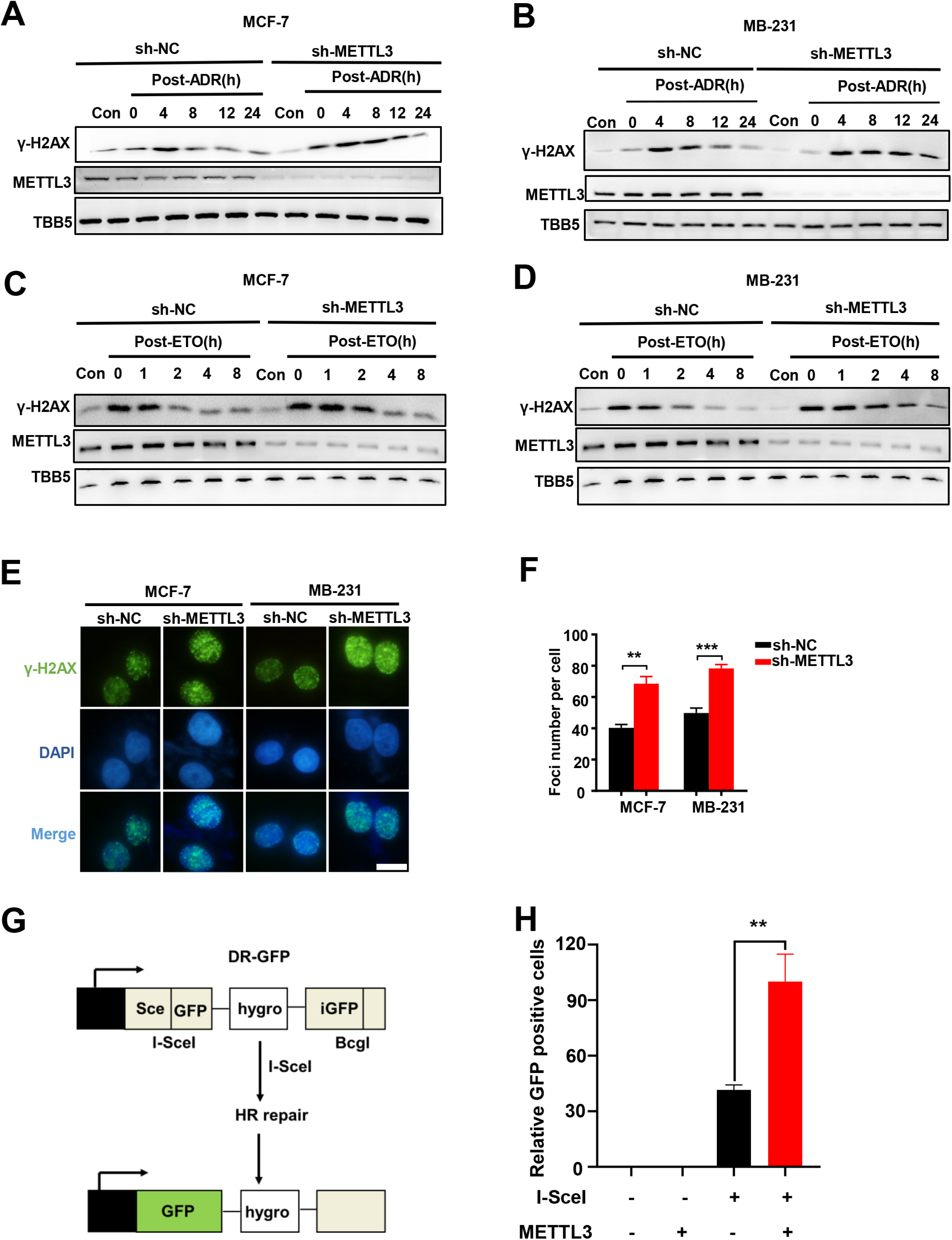
METTL3 regulates HR efficacy. (**A, B**) Western blot assay to determine γ-H2AX levels in control and METTL3-KD MCF-7 cells (**A**) or MB-231 cells (**B**) with ADR (0.5 μM) treatment for 1 hour following different recovery times. (**C, D**) Western blot assay to determine γ-H2AX levels in control and METTL3-KD MCF-7 cells (**C**) or MB-231 cells (**D**) with ETO (10 μM) treatment for 1 hour following different recovery times. (**E**) Immunofluorescence staining of γ-H2AX foci in different labeled cells with ADR treatment for 1 hour following 8 hours of recovery. (**F**) N=4 per group. (**H**) Relative quantification of the frequency of HR-mediated DSB repair in METTL3-ovexpressing U2OS cells. Data are represented as the mean ± SD of three biological repeats. N=3 per group. ^**^P< 0.01; ^***^ P< 0.001.

### EGF is the target of METTL3 and is regulated by m6A modification

A comprehensive assay combined with RNA-seq, MeRIP-qPCR, bioinformatics analysis, and literature retrieval was designed to explore the putative target(s) of METTL3-mediated m6A modification, which is involved in the regulation of both DNA repair and BC sensitivity to ADR (Fig. 3A). A total of 98 genes showed significant changes (p < 0.05; 42 up-regulated and 56 down-regulated) in METTL3-ovexpressing MCF-7 cells compared with control MCF-7 cells (Fig. 4B). Among these genes, 52 were shown to be modified by m^6^A in the exonic, 5’UTR, or 3’UTR of mRNA region in the m^6^A-Atlas, a comprehensive knowledgebase for unraveling the m^6^A epitranscriptome (Table 1) (Tang et al., 2021). Furthermore, literature retrieval identified 8 out of 52 genes that were reported to be involved in the regulation of DNA repair, among which EGF was highlighted because of its role in cancer progression, and DNA repair (Myllynen et al., 2011; Wilson et al., 2009; Yacoub et al., 2003). Next, we verified the expression of EGF regulated by METTL3. The mRNA levels of EGF increased in both METTL3-OV MCF-7 and MDA-MB231 cells compared with control cells (Fig. 3C, D), whereas knockdown of METTL3 down-regulated EGF expression (Supplementary Fig. S3A, B). Western blot analysis of whole cell lysates further verified the up-regulation of EGF by METTL3 (Fig. 3E, F and Supplementary Fig. S3C, D). Moreover, secreted EGF in the culture supernatants were examined by ELISA. The results indicated increased EGF levels in the medium of METTL3-OV MCF-7 and MB-231 cells (Fig. 3G, H), whereas down-regulated EGF was detected in the medium of METTL3-KD cells (Supplementary Fig. S3E, F). Furthermore, to validate the m^6^A modification in EGF mRNA, we performed a methylated RNA immunoprecipitation (meRIP)-qPCR assay using an m^6^A antibody followed by qPCR for the predicted region of the m^6^A sites. EGF mRNA exhibited the highest score by the sequence-based RNA adenosine methylation site predictor algorithm (Zhou et al., 2016). Using specific primers designed for the predicted m6A-harboring regions of EGF, the qPCR data showed that EGF was the target of METTL3 and regulated by METTL3-mediated m6A modification (Fig. 3I).

**Table 1.**
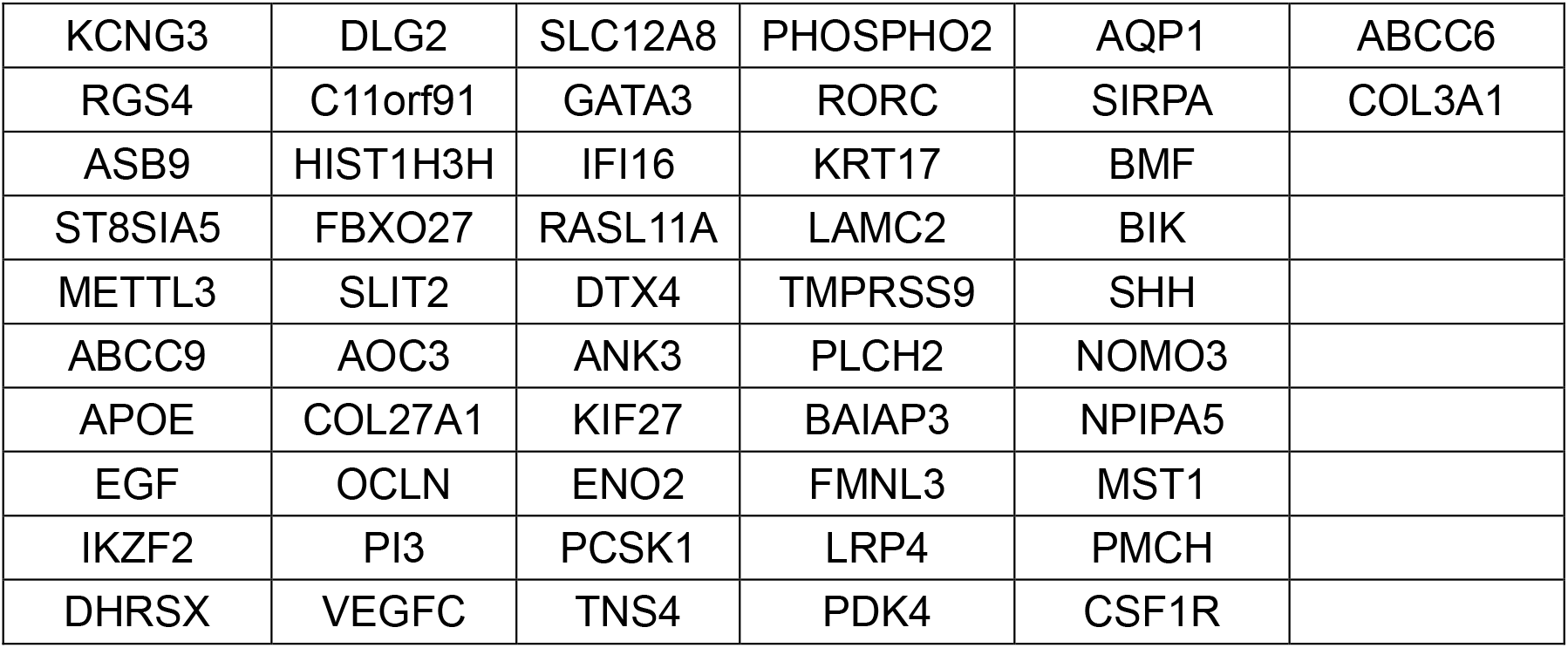
52 genes were showed to be modified by m^6^A in the exonic in this study.

**Table 2.**
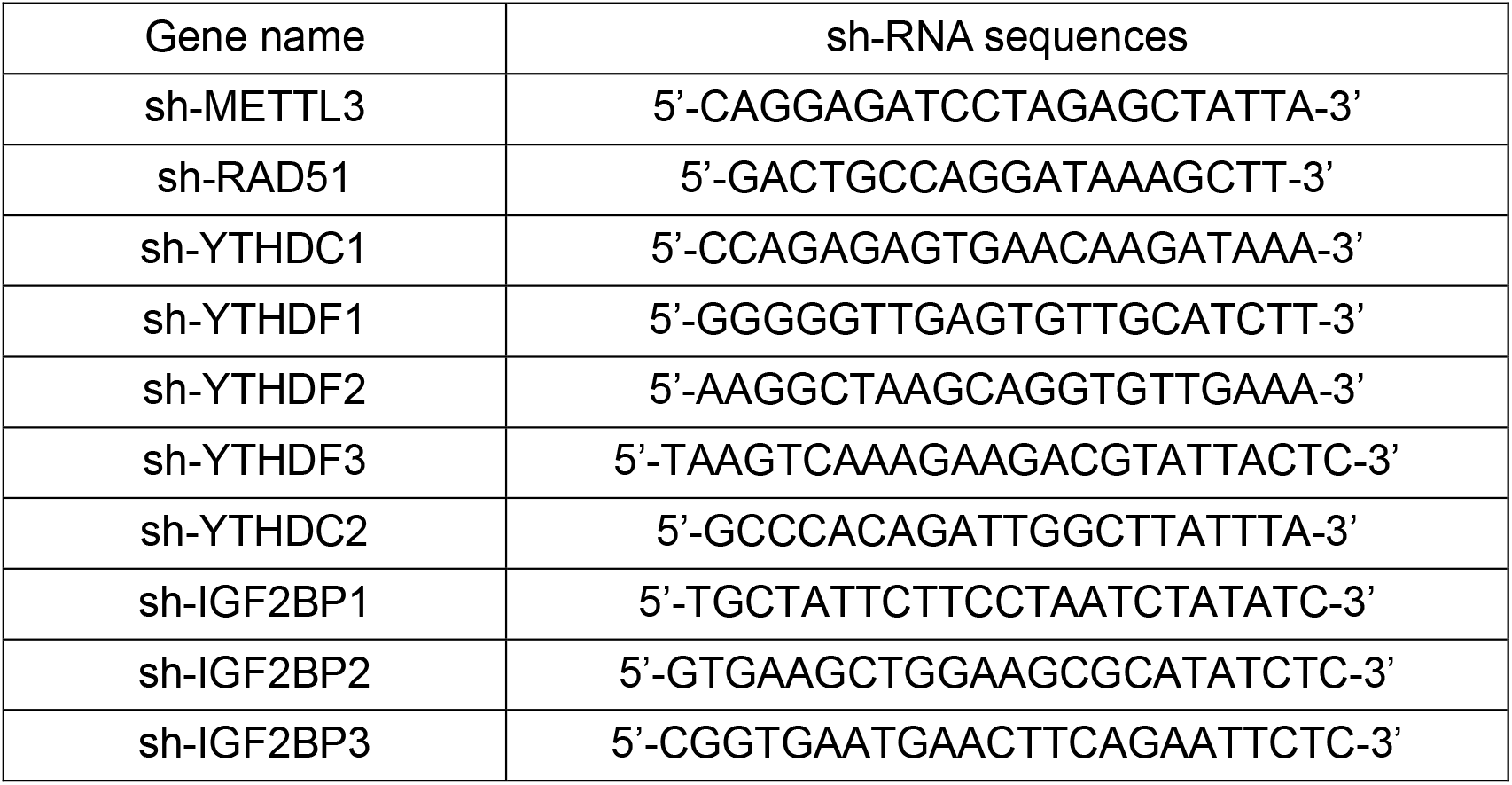
Sequences of the sh-RNA used in this study.

**Figure 3.**
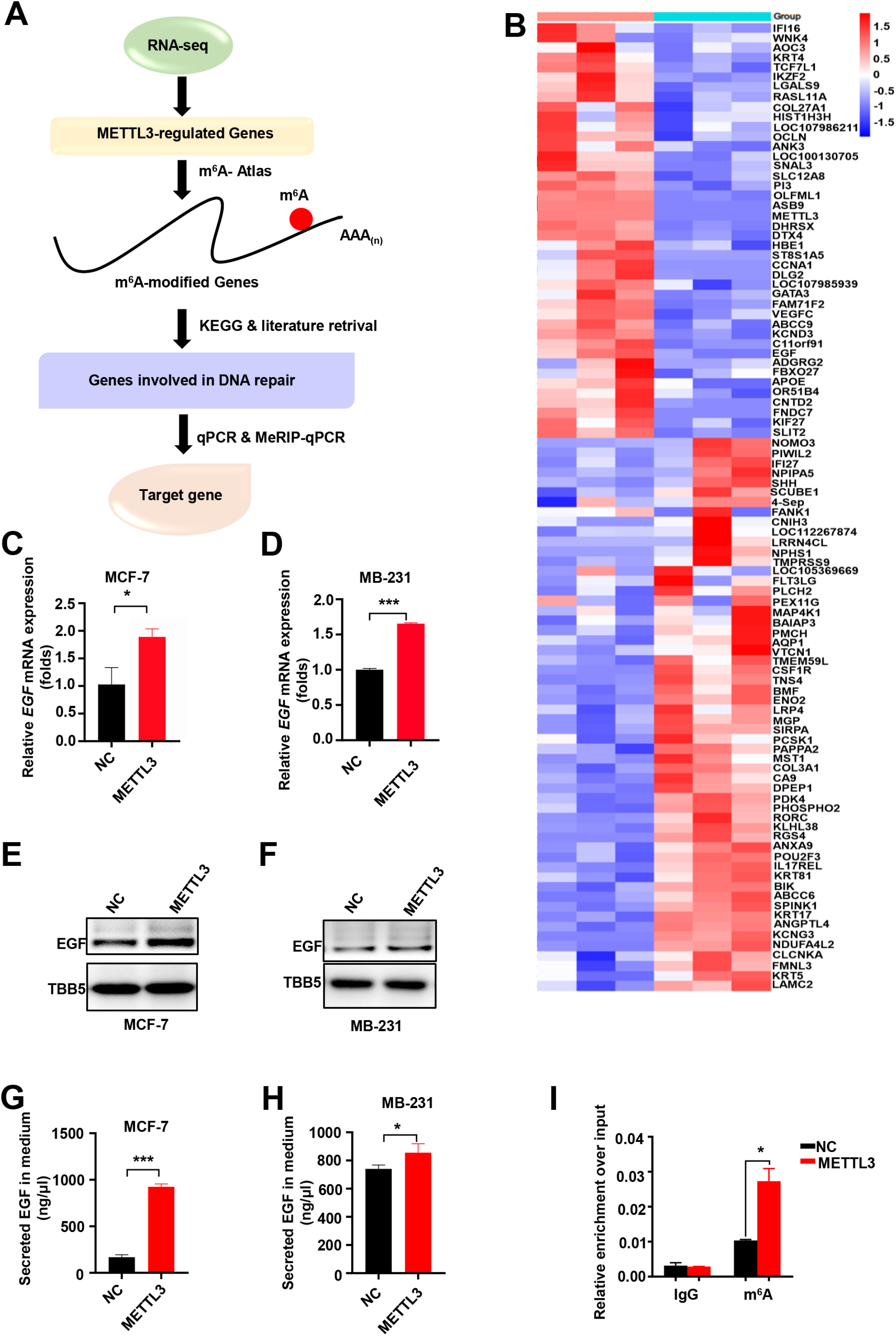
Identification of EGF as the targets of METTL3 in breast cancer. (**A**) Schematic of the screening progress of METTL3 targets in BC. (**B**) Heat map of RNA-seq to identify the genes regulated by METTL3 overexpression. (**C, D**) qRT-PCR was performed in METTL3-overexpressing MCF-7 (**C**) and MB-231 cells (**D**) to detect EGF expression. Data are expressed as the mean ± standard deviation (SD), n=3 per group. (**E, F**) Western blot analysis of EGF expression in METTL3-overexpressing MCF-7 (**E**) and MB-231 cells (**F**). (**G, H**) ELISA assay measuring secreted EGF in the medium of METTL3-overexpressing MCF-7 (**G**) and MB-231 cells (**H**). Data are expressed as the mean ± standard deviation (SD), n=3 per group. (**I**) MeRIP-qPCR analysis was used to assess the m^6^A levels of EGF mRNA in METTL3-overexpressing MCF-7 cells. The enrichment of m^6^A in each group was calculated by m^6^A-IP/input and IgG-IP/input. Data are expressed as the mean ± standard deviation (SD), n=2 per group. ^*^P< 0.05; ^***^ P< 0.001.

**Figure 4.**
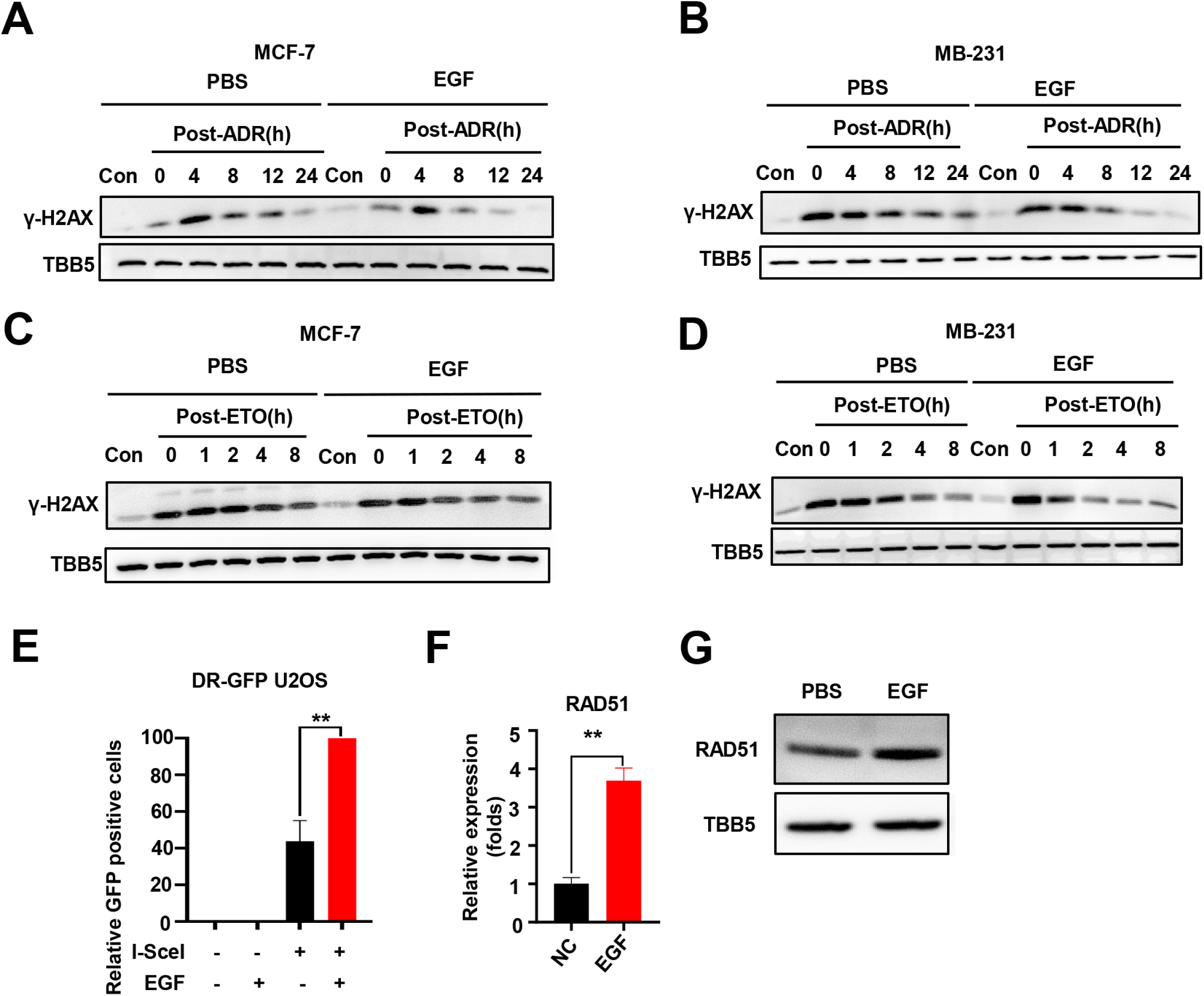
EGF promotes RAD51 expression and enhances HR. (A, B) EGF (10 ng/ml) enhanced DNA repair in MCF-7 (**A**) and MB-231 cells (**B**) treated with ADR (0.5 μM). (**C, D**) EGF enhanced DNA repair in MCF-7 (**C**) and MB-231 cells (**D**) treated with ETO (10 μM). (**E**) EGF treatment (10 ng/ml for 4 hours) enhanced HR efficacy in a GFP-based HR reporter system. Data are expressed as the mean ± standard deviation (SD), n=3 per group. (**F, G**) EGF augmented RAD51 mRNA (**F)** and protein (**G**) expression in MCF-7 cells. Data are expressed as the mean ± standard deviation (SD), n=2 per group. ^**^ P< 0.01.

### EGF regulates RAD51 expression and enhances HR activity

The EGF/EGFR signaling pathway has been reported to regulate DSB repair in lung cancer cells following X-irradiation by promoting both the NHEJ and HR pathways (Kriegs et al., 2010; Myllynen *et al*., 2011). Thus, we wondered whether EGF/EGFR is involved in METTL3-mediated DSB repair in BC cells treated with chemotherapeutic agents, such as ADR. First, we evaluated the effect of the EGF/EGFR signaling pathway on DSB repair in BC cells. MCF-7 and MB-231 cells were treated with ADR or ETO, respectively, and then released for different times with EGF treatment. Western blot analysis showed that the γH2AX levels markedly decreased in both ADR- and ETO-treated cells following EGF treatment, indicating that EGF enhanced DSB repair (Fig. 4A-D). Using a GFP-based HR reporter system, we confirmed the effect of the EGF/EGFR pathway on regulating HR activity (Fig. 4E and Supplementary Fig. S4A, B). To explore the molecular mechanism of EGF-mediated HR, we determined whether EGF/EGFR regulated the expression of core genes involved in HR, including *BRCA1, BRCA2, CtIP*, and *RAD51*. Our data showed that both EGF and METTL3 exhibited a slight effect on the regulation of BRCA1, BRCA2, and CtIP expression (Supplementary Fig. S4C, D). In contrast, the expression of RAD51 was markedly regulated by the EGF/EGFR signaling pathway (Fig. 4F, G; and Supplementary Fig. S4E-G).

### METTL3-modification of DNA repair is EGF/ RAD51 dependent

We next explored the effect of EGF/EGFR signaling on METTL3-mediated HR in BC cells. Western blotting analysis showed that knockdown of METTL3 down-regulated RAD51 expression, which was recovered by EGF treatment in BC cells (Fig. 5A, B). In contrast, overexpression of METTL3 up-regulated RAD51 expression, which was repressed by the EGFR inhibitors, erlotinib and gefitinib (Supplementary Fig. S5A, B). Both EGF and RAD51 levels were down-regulated in xenograft tissues derived from METTL3-KD MCF-7 cells (Fig. 5C). Accordingly, EGF treatment impeded METTL3-KD-mediated repression of DNA repair in both MCF-7 and MB-231 cells (Fig. 5D and E). The EGFR inhibitor, erlotinib, repressed DNA repair activities that were up-regulated by overexpression of METTL3 in both BC cells (Supplementary Fig. S5C, D). These data indicate that the EGF/EGFR signaling pathway regulates RAD51 expression and participates in METTL3-mediated HR.

**Fig. 5.**
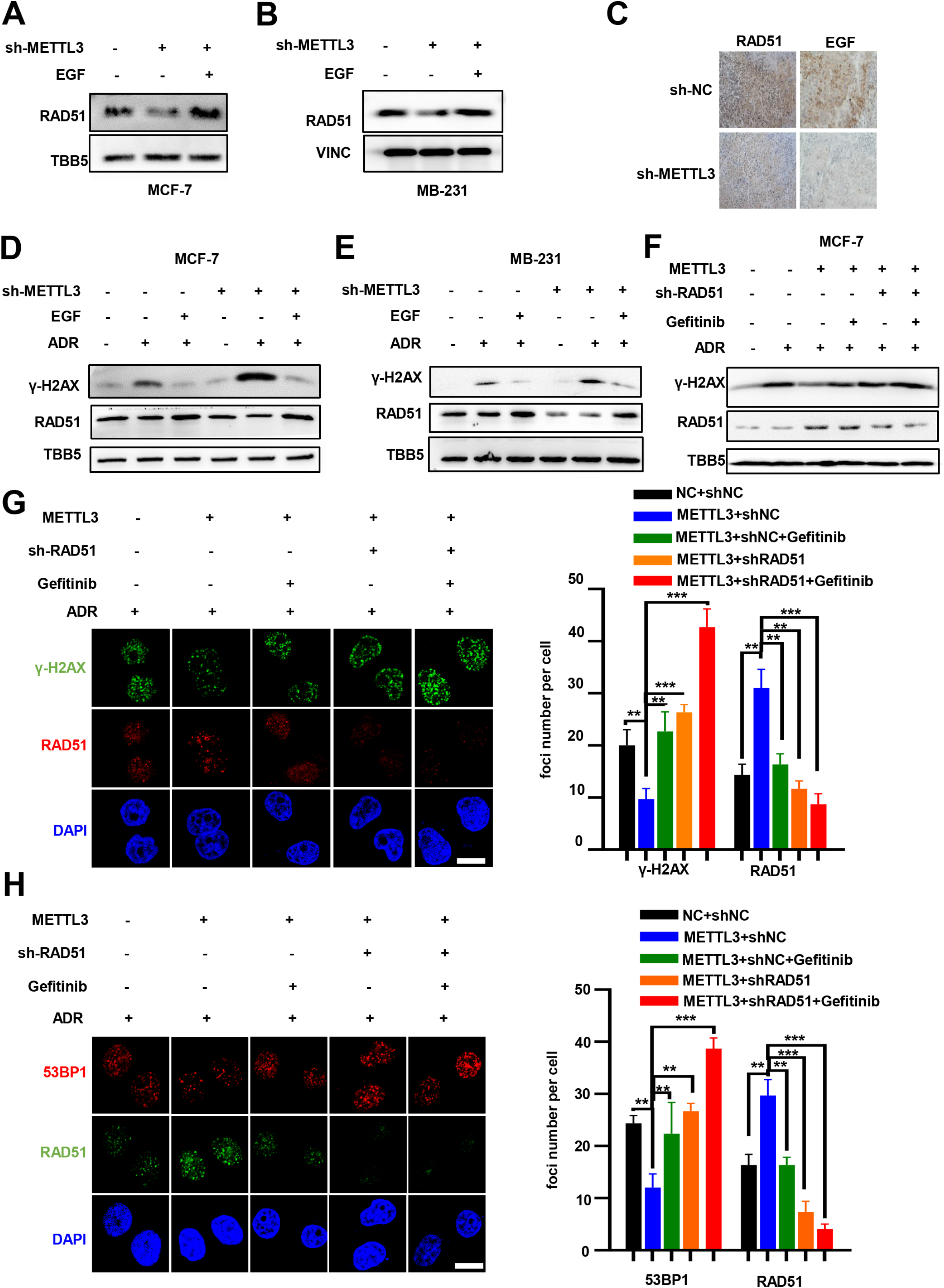
METTL3-modification of DNA repair is EGF/Rad51-dependent. (**A, B**) RAD51 protein levels in METTL3-KD MCF-7 (**A**) and MB-231 (**B**) cells treated with or without EGF. (**C**) Immunohistochemistry analysis of the expression of EGF and RAD51 in control and METTL3-KD tumor tissues. (**D, E**) WB analysis showing that treatment with 10 ng/ml EGF for 8 h restores DNA repair activity in METTL3-KD MCF-7 (**D**) and MB-231 (**E**) cells. (**F**) WB analysis showing that knocking down RAD51 or EGFR inhibitor Gefitinib (10 nM for 8 h) treatment in METTL3-OV cells decreases METTL3-enhanced DNA repair activity. (**G**) Immunofluorescence analysis of γ-H2AX and RAD51 foci in METTL3-OV cells ±RAD51 shRNA or gefitinib (10 nM) during ADR treatment (0.5 μM for 1 h and recovery for 8 h without ADR). The quantitative assay is on the right. N=3 per group. (**H**) Immunofluorescence analysis of 53BP1 and RAD51 foci in cells treated the same as in (**G**). The quantitative assay is on the right. N=3 per group. ^**^ P< 0.01; ^***^ P< 0.001.

We next investigated whether the effects of METTL3 on DNA repair were EGF/RAD51 dependent. As shown in Fig. 5F, the METTL3-OV MCF-7 cells were transfected with shRAD51 (shRNA of RAD51) for 36 hours, then treated with ADR for 1 hour and released for 8 hours with or without gefitinib/erlotinib treatment. Overexpression of METTL3 resulted in elevated DNA repair activity (shown by γH2AX down-regulation) which was reversed by treatment with siRAD51 or gefitinib/erlotinib (Fig. 5F and Supplementary Fig. S5E). Consistently, immunofluorescence analysis showed that the γH2AX and 53BP1 foci were down-regulated by overexpression of METTL3, which was reversed by treatment with siRAD51 or EGFR inhibitors (Fig. 5G and H). These results suggest that METTL3 augments HR in ADR-treated BC cells via the EGF/ RAD51 axis.

### YTHDC1 enhances the METTL3/m^6^A-regulated EGF/Rad51 axis

There are two major families of m^6^A “readers” that play a specific role in controlling the fate of methylated mRNA including the YTH family and the IGF2BP family (Deng *et al*., 2018; Wang *et al*., 2015; Xiao et al., 2016). To elucidate the specific m^6^A readers of EGF mRNA and to determine the m6A-dependent mechanism of EGF regulation, we performed qPCR assays to screen EGF-related m^6^A readers. Interestingly, knockdown of YTHDC1, but not other members of the YTH family or the IGF2BP family, down-regulated both EGF and RAD51 in MCF-7 cells (Fig. 6A and Supplementary Fig. S6A-C). Furthermore, knockdown of YTHDC1 reversed the METTL3-mediated up-regulation of EGF and RAD51 (Fig. 6B and C). Additionally, the direct interaction between *YTHDC1* and *EGF* transcripts was enhanced in METTL3-OV cells compared with that in control cells (Fig. 6D). The binding motif of YTHDC1 in EGF mRNA is UGG(m^6^A)CU, which is the preferentially binding motif of YTHDC1 (Xu et al., 2014). Furthermore, the YTHDC1 deficiency impaired the outcome of down-regulation of γH2AX foci in METTL3-OV MCF-7 cells (Fig. 6E). An MTT assay revealed that YTHDC1 depletion rendered MCF-7 cells more sensitive to ADR and reversed METTL3-induced ADR resistance in METTL3-overexpressing cells (Fig. 6F). These results were verified by morphological analysis (Fig. 6G). Taken together, our results suggest that YTHDC1 functions as an m6A “reader” to enhance EGF mRNA stability and augment HR and cell survival in ADR-treated BC cells.

**Fig. 6.**
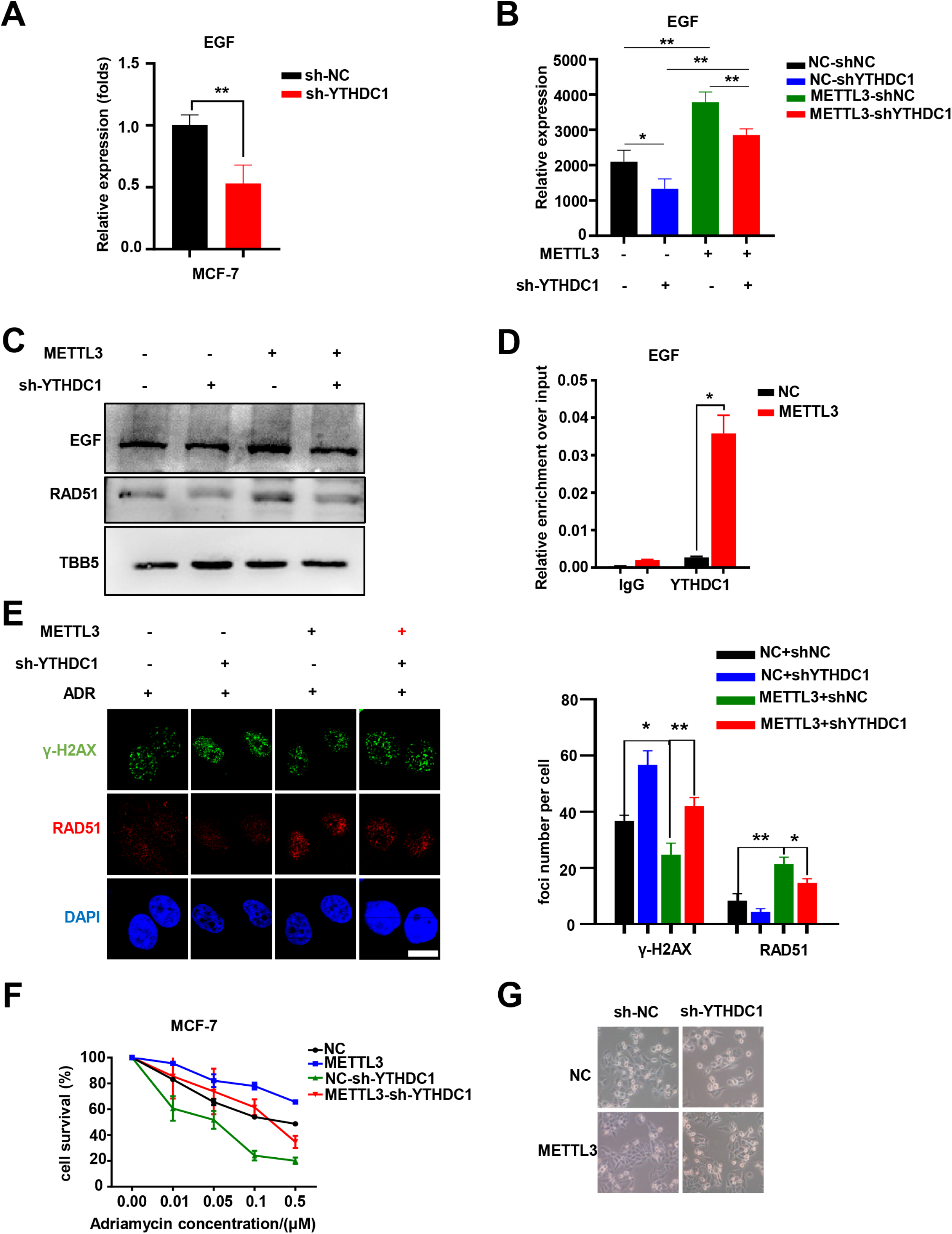
YTHDC1 is the “reader” of the METTL3/m6A-regulated EGF/Rad51 axis. (**A**) The expression of EGF in YTHDC1-silenced MCF-7 cells was detected by qRT-PCR. Data are expressed as the mean ± standard deviation (SD), n=3 per group. (**B**) The mRNA levels of EGF in control or METTL3-OV cells with or without knocking down YTHDC1. Data are expressed as the mean ± standard deviation (SD), n=3 per group. (**C**) WB assay determining the effect of YTHDC1 knockdown on EGF and RAD51 expression in control and METTL3-OV MCF-7 cells. (**D**) RIP-qPCR assay showing the enrichment of the EGF transcript in METTL3-OV cells. Data are expressed as the mean ± standard deviation (SD), n=2 per group. (**E**) Immunofluorescence analysis of γ-H2AX and RAD51 foci in METTL3-OV cells with knocked-down YTHDC1. The quantification of the foci numbers per cell are shown in the right panel. N=3 per group. (**F**) MTT assays were performed to detect the effect of YTHDC1 knockdown on ADR sensitivity in control and METTL3-OV MCF-7 cells. Data are expressed as the mean ± standard deviation (SD), n=3 per group. (**G**) Morphological analysis of control or METTL3-OV MCF-7 cells with or without YTHDC1 knockdown. Cells were treated with 0.5 μM ADR for 24 h. ^*^ P< 0.05; ^**^ P< 0.01; ^***^ P< 0.001.

## Discussion

A deficiency of DNA repair proteins is associated with carcinogenesis and elevated DNA repair activity contributes to drug resistance in cancer (Li *et al*., 2021; Lu *et al*., 2020). MELLT3 and its regulated m^6^A modification have been reported to be involved in DNA repair (Xiang *et al*., 2017; Zhang *et al*., 2020). However, the potential mechanism of METTL3 in DNA repair and chemotherapeutic response is poorly defined. In the present study, we demonstrated that m^6^A RNA methylation levels and METTL3 expression were elevated in BC cells. Biochemical and cell biological analyses revealed that inhibition of METTL3 sensitizes BC cells to ADR treatment with elevated DNA damage. Knockdown of METTL3 impaired HR activity and increased ADR-induced DNA damage through modification of the EGF/RAD51 axis. METTL3 promoted HR through m6A-dependent upregulation of EGF expression, which further augmented RAD51 expression. The m^6^A “reader,” YTHDC1, bound to the m^6^A-modified EGF transcript, protected EGF mRNA, and enhanced EGF expression (Fig. 7). This result was consistent with other studies showing that YTHDC1 is recruited to sites of DNA damage, bound m^6^A RNA, and increased the activity of DSB repair (Yu et al., 2021a; Zhang *et al*., 2020).

**Fig. 7.**
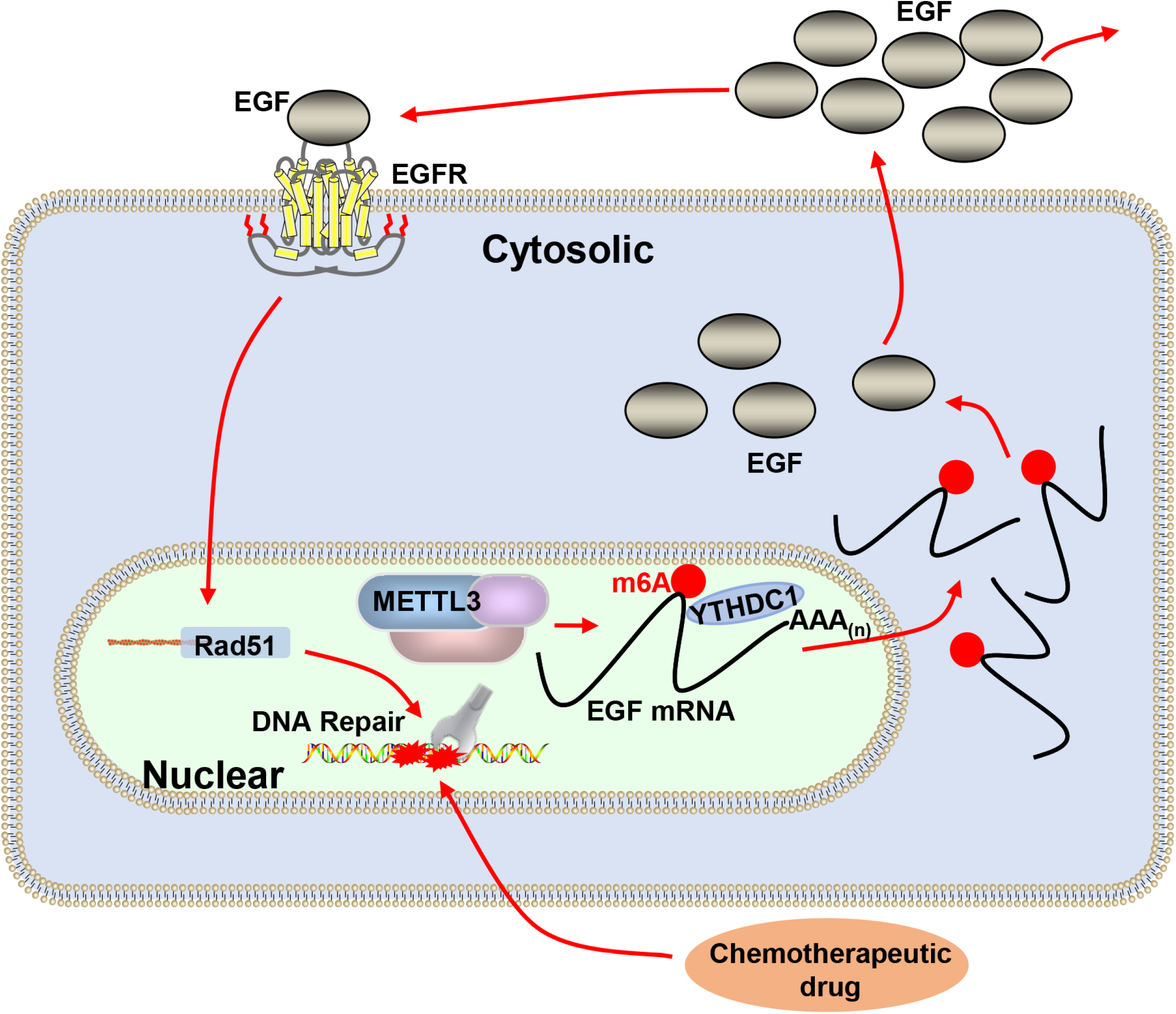
Proposed schematic diagram of the proposed mechanism elucidated in this study. METTL3 augments EGF transcript m6A modification, which was recognized by the reader YTHDC1, resulting increase of EGF expression. Elevated EGF promoted RAD51 expression, enhanced HR efficacy, and modified chemotherapeutic response of cancer cells.

Recently, RNA m^6^A modification and the core RNA methyltransferase, METTL3, were reported to play an important role in cancer chemotherapy (Deng *et al*., 2018). METTL3 was implicated in ADR resistance in MCF-7 cells by regulating the miR-221-3p/HIPK2/Che-1 axis (Pan *et al*., 2021). Our data further demonstrated that METTL3 knockdown markedly sensitized BC cells to ADR. ADR is one of the most effective antitumor agents for BC treatment, although it is limited by severe side effects. Various mechanisms have been proposed to explain ADR-induced cell death, including trapping topoisomerase II, formation of ADR-DNA adducts, and generation of free radicals that increase oxidative stress, which induce DNA damage and result in cell death. (Yang *et al*., 2014). We also showed that a deficiency in METTL3 inhibits DNA repair and increases the accumulation of DNA damage in ADR-treated BC cells, which further leads to cell death. A large number of studies have implicated dysfunctional DNA repair proteins, such as BRCA1, RAD51, FEN1, and Polβ, in BC initiation and progression (Li *et al*., 2021; Lu *et al*., 2020; Martin et al., 2007; Thacker, 2005; Wang et al., 2019; Wang et al., 2020b; Xia et al., 2021). Elevated DNA repair activity contributes to drug resistance and limits the efficacy of chemotherapeutic agents (Long et al., 2021; Lu *et al*., 2020). RNA m^6^A modification has been reported to be involved in the regulation of DNA repair. Xiang *et al*. reported that METTL3-mediated m^6^A RNA accumulates at UV-damaged sites in DNA, which further recruits DNA polymerase κ (Pol κ) to damaged sites to facilitate NER and cell survival (Xiang *et al*., 2017). Zhang et al. reported METTL3-m6A-YTHDC1 mediated HR in U2OS cells exposed to X-rays or treatment with Zeocin (Zhang *et al*., 2020). Another recent study found that METTL3-METTL14 is active in vitro on double-stranded DNA containing a cyclopyrimidine dimer, and the m6A reader, YTHDC1, is recruited to sites of DNA damage (Yu *et al*., 2021a). Our data demonstrated that silencing METTL3 inhibited the repair of DSBs induced by ADR. Nevertheless, the molecular mechanism of DNA repair regulation by METTL3/m6A remains undefined.

There are two major pathways to repair DSBs including error-free HR and error-prone NHEJ (Sonoda *et al*., 2006). Using a GFP-based reporter system, we found that METTL3 regulated HR, but not NHEJ in DSB repair, which is consistent with the study of Zhang et al (Zhang *et al*., 2020). However, Gene Ontology (GO) analysis of METTL3-associated gene expression profiles based on RNA-seq in MCF-7 cells did not identify enriched “DNA damage” or “DNA repair” categories (data not shown). This data is supported by Xiang et al. showing that GO analysis of METTL3-dependent, UV-induced methylated transcripts only identified “DNA damage” as weakly enriched (Xiang *et al*., 2017). Zhang et al. also reported that METTL3 does not change active RNA polymerase II binding to DNA lesions and has no effect on RNA transcription at DSBs. Another study showed that METTL3 catalyzes m6A modification of FEN1 mRNA, which is recognized and stabilized by IGF2BP2 in hepatocellular carcinoma cells. (Pu et al., 2020). However, our RNA-seq data showed no change in FEN1 expression with METTL3 overexpression in MCF-7 cells, suggesting that METTL3-mediated FEN1 expression may be cell specific.

EGF/EGFR signaling is important to many biological processes, such as cell proliferation, cell division, and tissue development. Human cancer tissues express high levels of growth factors and their receptors, such as EGF/EGFR, which exhibit autocrine or paracrine regulation of cancer growth (Knowlden et al., 2003; Mendelsohn and Baselga, 2000; Singh and Harris, 2005). Aberrant activation of EGF/EGFR signaling contributes to cancer proliferation, epithelial-mesenchymal transition, and metastasis (Mendelsohn and Baselga, 2000). Therefore, targeting EGFR by antibodies or small molecule tyrosine kinase inhibitors has been used successfully to treat various malignancies, including BC (Barzegar et al., 2017; Ueno and Zhang, 2011). Based on RNA-seq, MeRIP-qPCR, and the screening strategy, we determined that EGF is regulated in a METTL3-m6A-YTHDC1-dependent manner in BC cells. Overexpression of METTL3 markedly increased EGF expression and secretion, whereas knockdown of METTL3 resulted in the opposite effect. We further demonstrated that EGF plays an important role in METTL3-mediated HR in BC cells during ADR treatment. Our results are consistent with previous reports showing that EGF/EGFR signaling plays a role in DSB repair (Myllynen *et al*., 2011). Moreover, dysregulated EGF/EGFR signaling modulates the expression of several genes involved in DNA repair including ERCC1, XRCC1, RAD51, and RAD50 (Kryeziu et al., 2013; Yacoub *et al*., 2003). Accordingly, we found that EGF/EGFR signaling regulated RAD51 expression in BC cells and the regulation of RAD51 expression by METTL3 was EGF-dependent. Furthermore, knockdown of RAD51 or inhibition of EGFR by gefitinib/erlotinib impaired the effect of METTL3 on the up-regulation of DNA repair activity. Although other studies and our RNA-seq data suggest that there may be other factors involved in the regulation of METTL3-mediated DNA repair and response to chemotherapy (Cai *et al*., 2018; Pan *et al*., 2021; Wang *et al*., 2020a), our experiments demonstrate a role for EGF in the regulation of HR activity and ADR sensitivity in BC.

The m^6^A modification of EGF mRNA has been detected in human kidney and embryonic stem cells by different groups using m^6^A-REF-seq (GSE125240) and MAZTER-seq (GSE122961), respectively (Garcia-Campos et al., 2019; Zhang et al., 2019). Our MeRIP-qPCR assay identified that m^6^A modification of EGF mRNA was augmented in METTL3-expressing BC cells compared with that in control cells. The m^6^A-modified EGF mRNA was recognized and bound by YTHDC1, which further promoted EGF synthesis and secretion. We demonstrated that the regulation of EGF expression by METTL3 was m^6^A-YTHDC1-dependent and knockdown of YTHDC1 impeded the up-regulation of EGF in METTL3-overexpressing cells, and restored the sensitivity of MCF-7 cells to ADR. Moreover, knockdown of YTHDC1 impeded METTL3-enhanced RAD51 expression and inhibited recruitment of RAD51 to damaged sites. Our data combined with the studies of Zhang et al. and Yu et al. demonstrate that YTHDC1 plays an important role in DNA repair (Yu *et al*., 2021a; Zhang *et al*., 2020). Our results further suggest the involvement of YTHDC1 in the regulation of EGF mRNA stability. Although YTHDC1 is primary known as a nuclear m^6^A reader, which regulates mRNA splicing through the recruitment and modulation of pre-mRNA splicing factors such as SRSF3 (Xiao *et al*., 2016), our data suggest that YTHDC1 may contribute to the stability of cytosolic mRNA, including EGF, through m^6^A. Our hypothesis is supported by previous studies suggesting that YTHDC1 can shuttle between the nucleus and the cytoplasm (Rafalska et al., 2004), to process mature mRNAs, including MAT2A (Shima et al., 2017). Moreover, Roundtree et al. showed that YTHDC1 mediates the export of methylated mRNA from the nucleus to the cytoplasm, resulting in nuclear clearance of mRNAs and accompanying their resulting cytoplasmic abundance (Roundtree et al., 2017). These results suggest multiple processes through which YTHDC1 regulates the processing of mature mRNA, whereas the detailed molecular mechanism warrants further study.

Collectively, we showed an effect of METTL3 on HR via the m6A-YTHDC1-dependent regulation of the EGF/RAD51 axis, and demonstrated a role for METTL3 in the response of BC chemotherapy. Our results suggest that the development of METTL3 inhibitors or targeting its pathway may lead to promising treatments for cancer patients.

## Materials and methods

### Plasmid construction

The oligonucleotide 5′-CAGGAGATCCTAGAGCTATTA-3′ was used for construction of METTL3-KD lentivirus vectors as previously described (Xiang *et al*., 2017). The lentivirus vectors were constructed and purified by the Corues Biotechnology Company (Nanjing, China). For knockdown of RAD51, YTHDC1 and other “readers,” and the silencing plasmids containing shRNA sequences were constructed based on psilencer3.0-H1. The shRNA sequences are listed in Supplementary Table 2. All plasmids were verified by sequencing.

### Cell culture and the development of stable cell lines

MCF-7 and MDA-MB-231 (MB-231) were purchased from the Shanghai Institute of Cell Biology, Chinese Academy of Science. Cells were cultured in the recommended medium supplemented with 10% fetal bovine serum (FBS, Invigentech), 1% penicillin, and 1% streptomycin, and incubated in an incubator with 5% CO_2_ at 37 °C. For METTL3-overexpressing (OV) or – knockdown (KD) MCF-7 and MB-231 stable cells, the cells were infected with specific lentivirus vectors for 48 h and then selected with puromycin for two weeks. All cell lines were confirmed to be negative for mycoplasma contamination.

### M^6^A Dot blotting

Total RNA was isolated using the Trizol method and mRNAs were isolated with the GenElute™ mRNA Miniprep Kit (Sigma). The concentration and purity of the mRNA were measured using a NanoDrop 2000. The mRNAs were denatured by heating to 95°C for 5 min, followed by chilling on ice. Next, the mRNAs (50∼100ng) were spotted directly onto a positively-charged nylon membrane (GE Healthcare, USA) and air-dried at room temperature for 5 min. The membrane was then ultraviolet (UV) crosslinked using a Ultraviolet Crosslinker, washed with PBST for 5 minutes, blocked with 5% nonfat milk in TBST, and then incubated with anti-m6A antibody (A17924, ABclonal) overnight at 4°C. HRP-conjugated anti-rabbit IgG secondary antibody was added to the membrane for 1.5 h at room temperature with gentle shaking, followed by development with enhanced chemiluminescence. Last, 0.02% methylene blue staining was used to verify that equal amounts of mRNA were spotted onto the membrane.

### Drug sensitivity assay

Cells were seeded into 96-well plates at 3000 cells per well for at least three parallel experiments. Twenty-four hours later, cells were exposed to ADR at increasing concentrations (0, 0.01, 0.05, 0.1, 0.5, and 1 μM) for 48 h; 5-FU at increasing concentrations (0, 0.1, 0.5, 1, 5 μM) for 48 h; carboplatin at increasing concentrations (0, 10, 50, 100, 500 μM) for 48 h; paclitaxel at increasing concentrations (0, 0.01, 0.1, 1 nM) for 48 h; or DDP at increasing concentrations (0, 2, 4, 6, 8 μM) for 48 h. Chemotherapeutic drug-treated cells were incubated with 10 μL 3-(4,5)-dimethylthiazol (-z-y1)-3,5-diphenyltetrazolium bromide (MTT) solution (5 mg/mL, Sigma-Aldrich, St Louis, MO, USA) for 4 h. The media was replaced with 100 μL dimethyl sulfoxide (DMSO, Sigma-Aldrich) to dissolve the formazan crystals within 10 min.

### RNA-seq and analyses

RNA-Seq was performed by oeBiotech Inc. (Shanghai, China). For RNA sequencing, purified RNA from METTL3 overexpressing or control cells was used for library construction with the Illumina TruSeq RNA Sample Prep Kit (FC-122-1001) and then sequenced with an Illumina HiSeq 2000. Raw reads were aligned to the human genome, GRCh37/hg19, by Bowtie2. Differentially expressed genes (DEGs) between METTL3-OV and the control samples were identified using the limma-voom method. A heatmap clustered by k-means was used to show DEGs or transcripts. The raw sequencing data were deposited in the Gene Expression Omnibus database (accession to cite for these SRA data: PRJNA743152).

### RNA immunoprecipitation (RIP)

RNA immunoprecipitation (RIP) assays were conducted using the EZ-Magna RIP™ RNA-Binding Protein Immunoprecipitation Kit (Merck Chemicals (Shanghai Co., Ltd). The anti-YTHDC1 antibody for RIP was purchased from Cell Signaling Technology, Inc. (# 81504S).

### M^6^A-RNA immunoprecipitation (MeRIP) and MeRIP-qPCR

M6A enrichment followed by qRT-PCR was used to quantify the changes in m6A methylation of the target gene using the Magna MeRIP m6A Kit (Millipore, MA) following the manufacturer’s instructions. Briefly, 5 μg of fragmented mRNA extracted from MCF-7 stable cells was incubated with 5 μg of m6A antibody (A17924, ABclonal). Methylated mRNA was eluted by free m6A from the beads and purified with the GenElute™ mRNA Miniprep Kit (MRN70, Sigma). One tenth of the fragmented RNA was saved as an input control for standardization. The relevant enrichment of m^6^A from METTL3 in each sample was analyzed by RT-qPCR.

### Immunofluorescence

For immunofluorescence assays, cells were washed with PBS for three times then fixed with 4% formaldehyde for 10 min at room temperature. After permeabilization with 0.1%Triton X-100 for 10 min, cells were blocked with 3% BSA for 1 h. Then, cells were incubated with indicated primary antibodies overnight at 4 °C. Following washed with PBST for three times, cells were incubated with fluorescent secondary antibodies for 2 h at room temperature. Subsequently, cells were stained with DAPI and visualized under a fluorescence microscope (Nikon 80I 10-1500×).

### Immunohistochemistry

Immunohistochemical staining was performed as previously described (Lu *et al*., 2020). Briefly, tumor tissues were fixed in 4% polysorbate. Paraffin-embedded sections from tissue specimens were deparaffinized and heated at 100°C in 10 mM citrate buffer (pH 6.0) for 15 min for antigen retrieval. Slides were incubated with primary antibody at 4 °C overnight, followed by incubation with secondary antibody at room temperature and visualized using a DAB Kit (Bioworld). Then, it was redacted with hematoxylin. The expression levels of target proteins in tissue were examed according to the semiquantitative immunoreactivity score (IRS).

### Apoptosis assay

METTL3-KD or control MCF-7 or MB-231 cells were treated with doxorubicin for 24 hours and replaced with fresh media. After other 24 hours recovery, about 1×10^5^ cells per well were collected and stained with both Annexin V and propidium iodide (PI). Apoptosis was analyzed by flow cytometry using the BD FACSverse.

### ELISA

The cell culture media were centrifuged at the speed of 5000r.p.m for 5 min and supernatant were collected for EGF measurement using commercial kits (SenBeiJia Biological Technology Co., Nanjing, China) according to manufacturer’s instructions.

### Statistical analysis

Statistical analysis was performed with GraphPad Prism 8.0. Statistical significance was determined using a two-tailed Student’s t-test or analysis of variance in the case of comparisons among multiple groups. P <0.05 was considered statistically significant.

Additional materials and methods are described in the Supplementary Methods.

## Acknowledgements

This work was supported by National Natural Science Foundation of China (81872284), Natural Science Fund of Colleges and Universities in Jiangsu Province (19KJA180010) and the Priority Academic Program Development of Jiangsu Higher Education Institutions.

## Author contributions

E.L., M.X., Y.D., and K.L. performed study concept and design; Z.G., Z.H., K.L., L.H., and F.P. performed development of methodology and writing, review and revision of the paper; E.L., M.X., Y.D., F.J., Z.H., K.L. and Z.G. provided acquisition, analysis and interpretation of data, and statistical analysis; F.P., Z.H. and Z.G. provided technical and material support. All authors read and approved the final paper.

## Competing interests

There are no potential conflicts of interest to disclose.

---

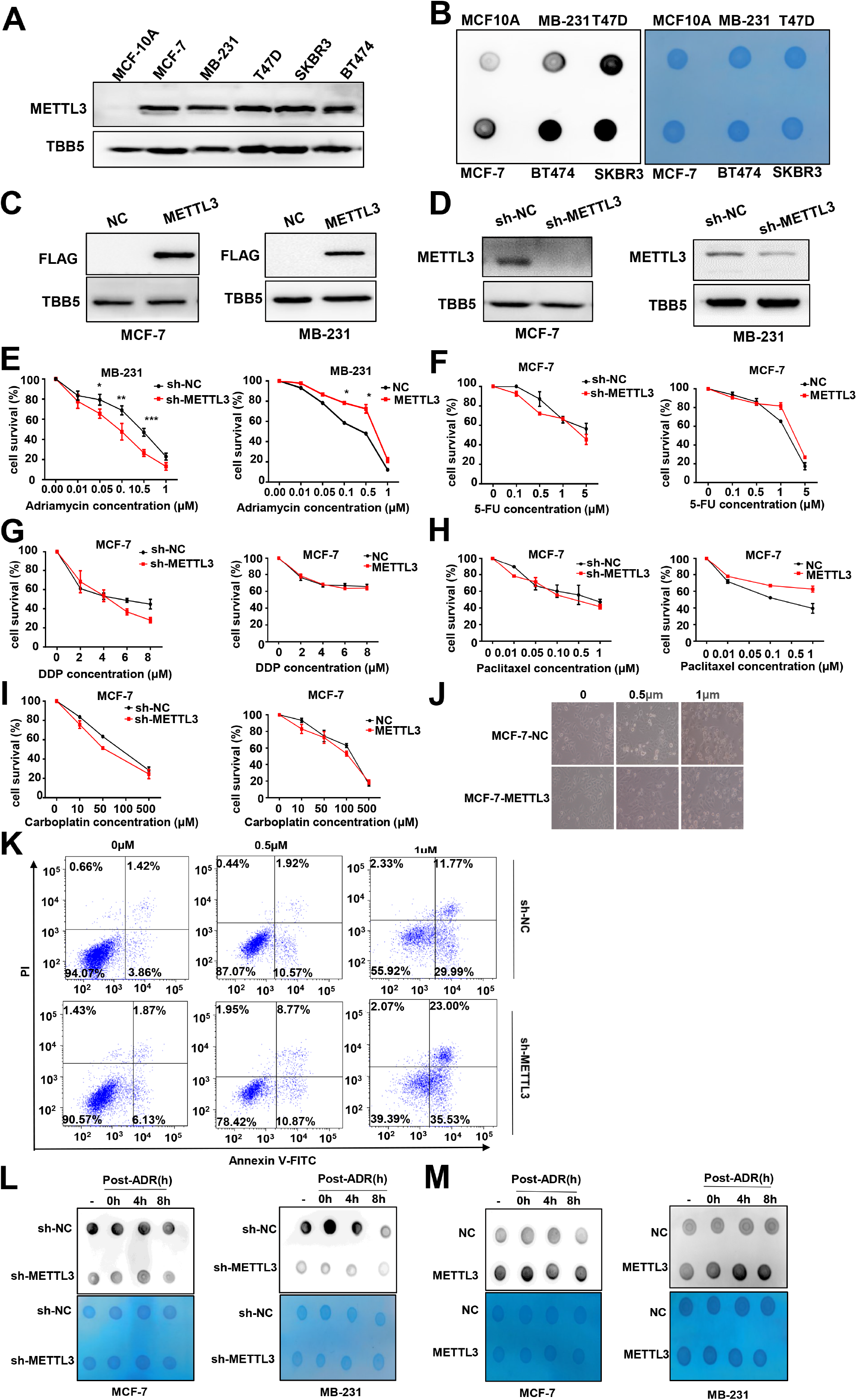

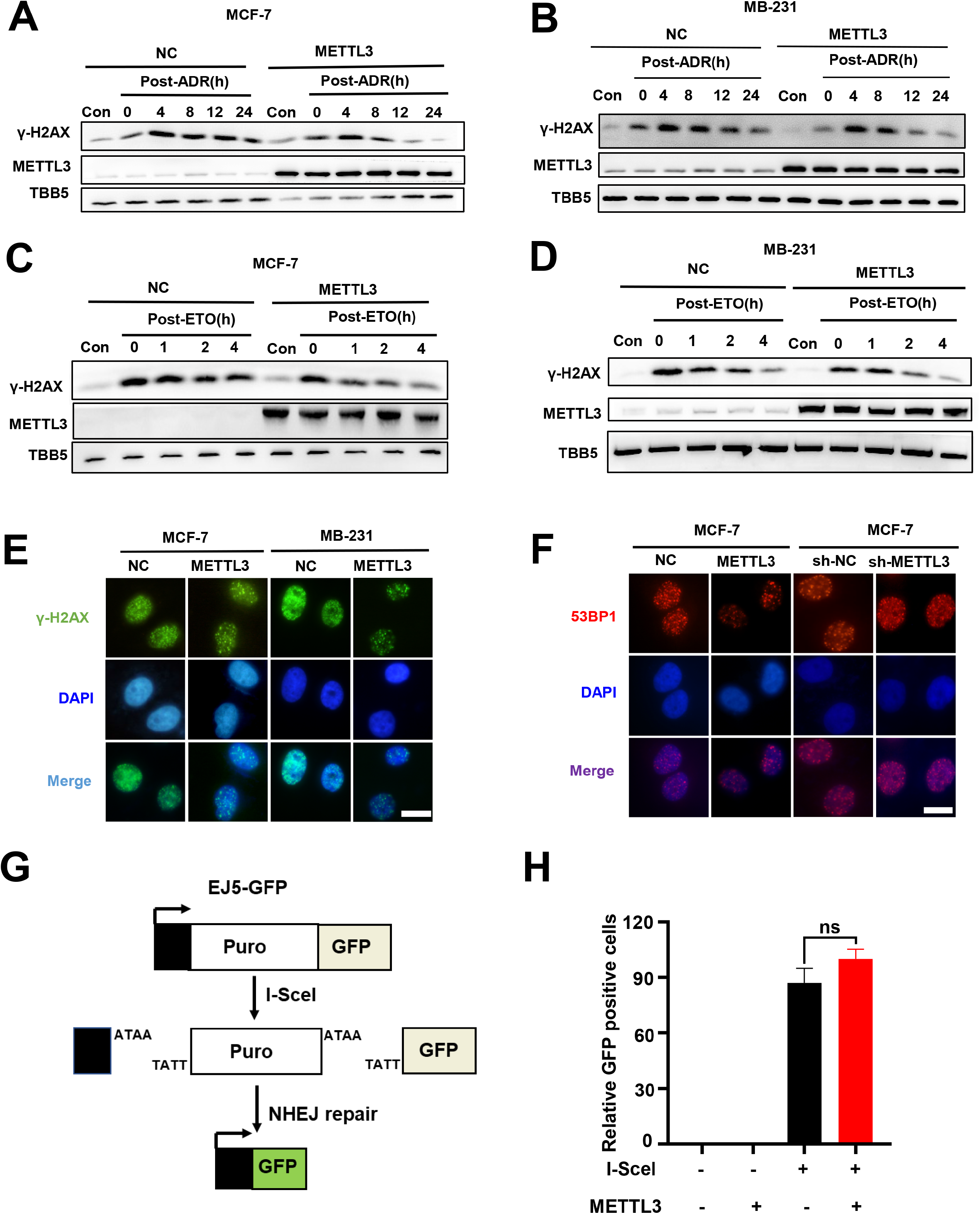

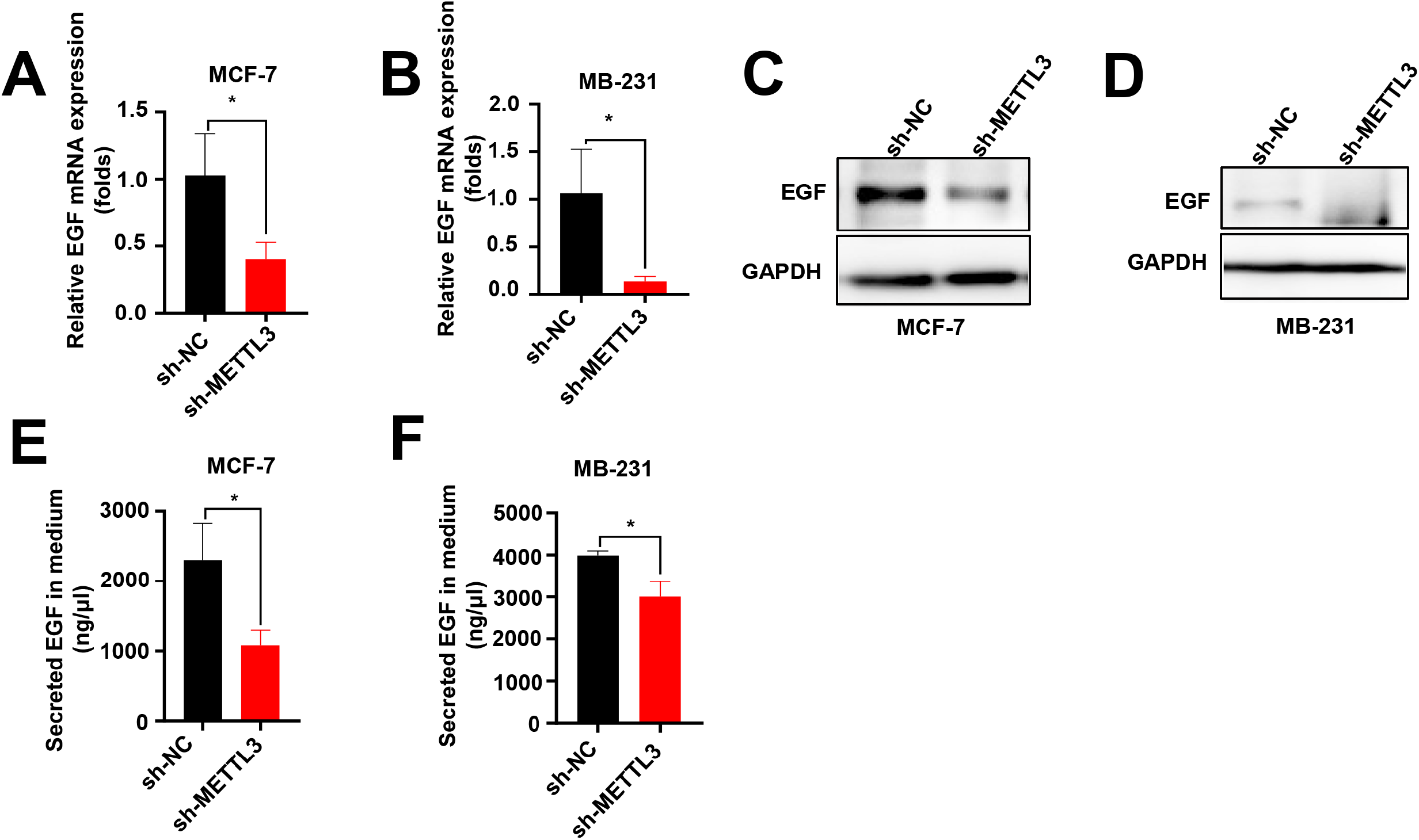

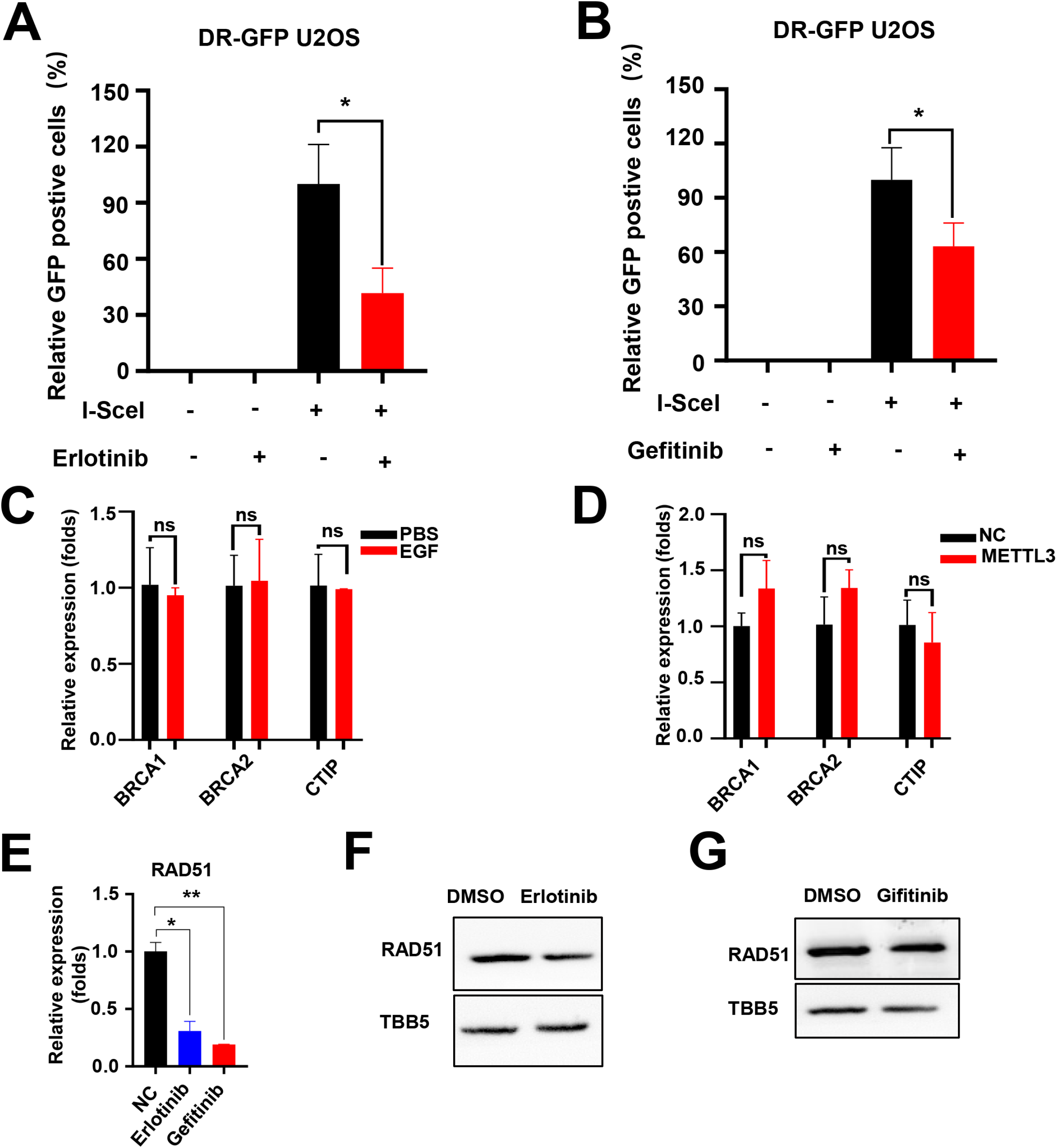

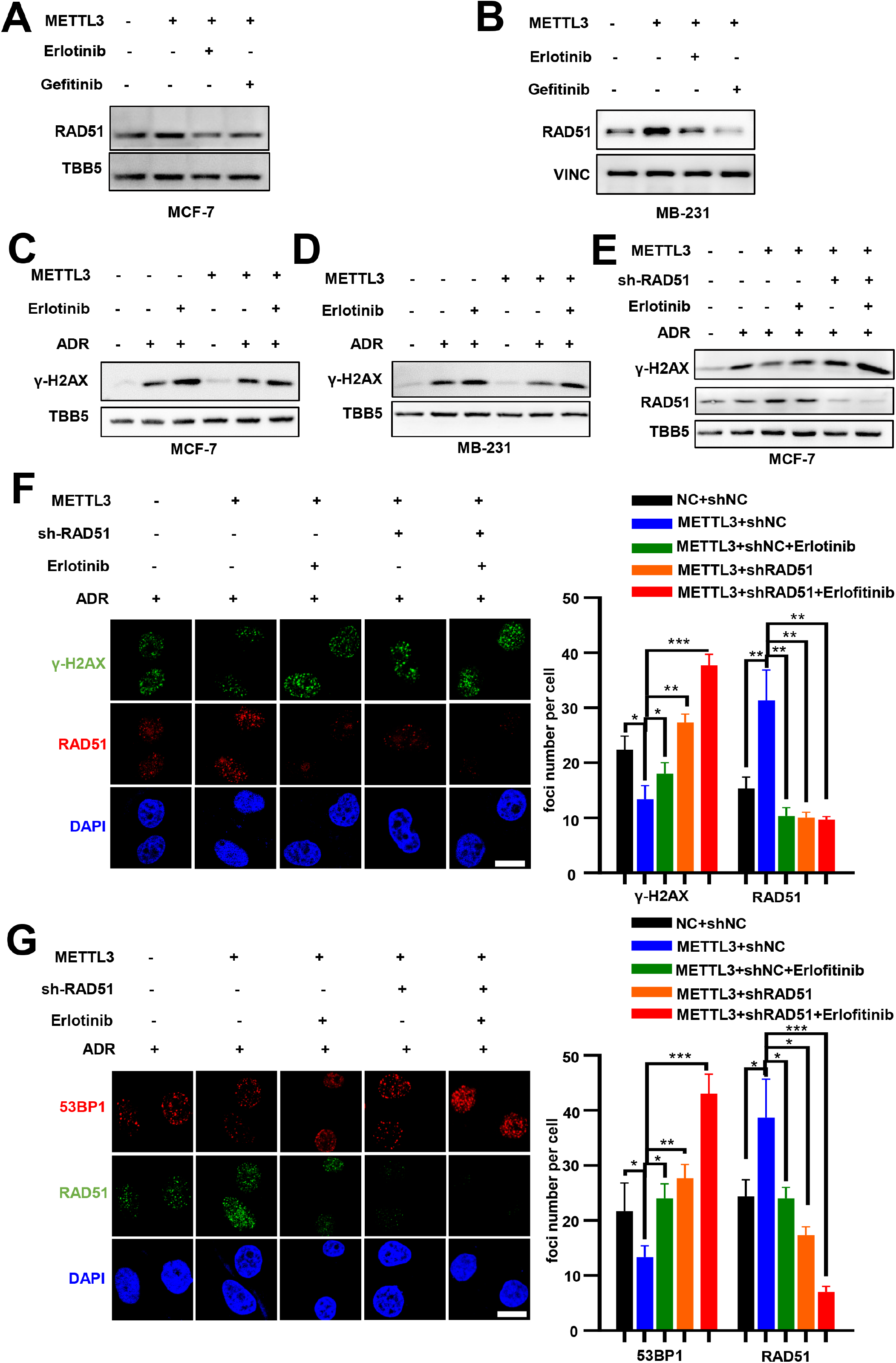

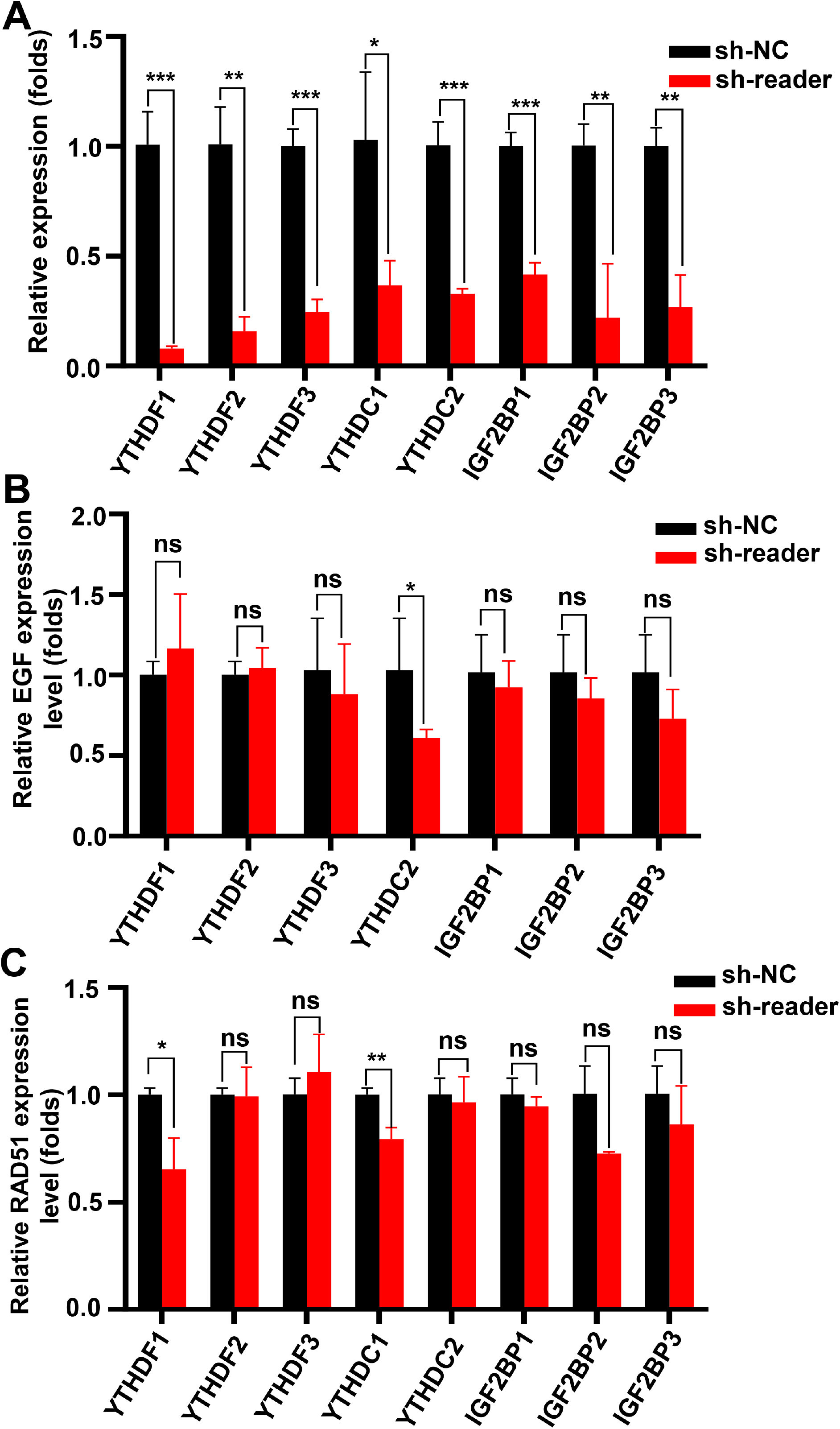

## Notes

### Competing Interest Statement

The authors have declared no competing interest.

